# Assessing metacommunity processes through signatures in spatiotemporal turnover of community composition

**DOI:** 10.1101/480335

**Authors:** Franck Jabot, Fabien Laroche, François Massol, Florent Arthaud, Julie Crabot, Maxime Dubart, Simon Blanchet, François Munoz, Patrice David, Thibault Datry

**Author notes:** Corresponding author: Franck Jabot. **Authorship statement:** all authors conceived the original idea and gave final approval for publication. FJ, FA, JC and MD performed the analyses. FJ lead the writing of the draft with contributions from all authors. **Data accessibility statement:** all data are available online – see l. 673-678.

## Abstract

Although metacommunity ecology has been a major field of research in the last decades, with both conceptual and empirical outputs, the analysis of the temporal dynamics of metacommunities has only emerged recently and consists mostly of repeated static analyses. Here, we propose a novel analytical framework to assess metacommunity processes using path analyses of spatial and temporal diversity turnovers. We detail the principles and practical aspects of this framework and apply it to simulated datasets to illustrate its ability to decipher the respective contributions of entangled drivers of metacommunity dynamics. We then apply it to four empirical datasets. Empirical results support the view that metacommunity dynamics may be generally shaped by multiple ecological processes acting in concert, with environmental filtering being variable across both space and time. These results reinforce our call to go beyond static analyses of metacommunities that are blind to the temporal part of environmental variability.

## Introduction

A main goal of community ecology is to understand the determinants of species diversity at different spatial scales. Metacommunity theory has emerged as a framework to investigate the spatial distribution of species and the dynamics of spatially structured ecosystems (Leibold et al. 2004, Massol et al. 2011, Guichard 2017). Metacommunity theory has been originally proposed to include four main (not mutually exclusive) paradigms explaining the coexistence of species (Leibold et al. 2004, Shoemaker & Melbourne 2016, Fournier et al. 2017; but see also criticism of Brown et al. 2017), and synthesizing how basic processes can drive metacommunity assembly. The patch-dynamic paradigm focuses on the processes of competition, colonization and extinction in networks of patches (Hastings 1980, Tilman 1994, Calcagno et al. 2006). The species-sorting paradigm focuses on the differential responses of species to environmental heterogeneity across the landscape to explain large-scale and local coexistence as the result of environmental filters and local adaptation (Chase & Leibold 2003). The mass-effect paradigm focuses on source-sink dynamics among communities, with species potentially co-occurring in patches where they are maladapted due to the influx of dispersing individuals (Amarasekare & Nisbet 2001, Mouquet & Loreau 2003). The neutral paradigm solely considers the effects of demographic stochasticity and dispersal limitation on community dynamics, thus explaining local species co-occurrence as a stochastic process driven by species frequencies at a larger scale and immigration (Hubbell 2001). These four simplistic views of metacommunities were defined to encompass the main models and assumptions on coexistence mechanisms, both in theory and in empirical studies (Cottenie 2005, Shoemaker & Melbourne 2016, Ulrich et al. 2017).

Metacommunity paradigms and related models have mostly been used to analyse spatial patterns of metacommunity composition at a single date, assuming that metacommunities are then at a dynamical equilibrium (Logue et al. 2011, Heino et al. 2015), and often with statistical models not founded on dynamical models (but see Azaele et al. 2006). Specifically, when spatial environmental variation is hypothesized to play a role, the most common approach is to perform variance partitioning (Borcard et al. 1992, Cottenie 2005, but see e.g., Leibold and Mikkelson 2002, Ulrich et al. 2017). It consists in partitioning the observed spatial variation of community composition into spatial and environmental components (Borcard et al. 1992, Cottenie 2005, Peres-Neto et al. 2006). The effect of the spatial component is then expected to reflect a combined effect of dispersal and ecological drift, while the effect of the environmental component should summarize differential species responses to environmental variation. Such analyses of static spatial patterns of metacommunities provided insights on the processes structuring metacommunities across biomes, taxa and along environmental gradients (Cottenie 2005, Henriques-Silva et al. 2013, Heino et al. 2015). However, results on simulated datasets suggest that partitioning alone does not allow unambiguous inference of metacommunity dynamics (Gilbert & Bennett 2010, Peres-Neto & Legendre 2010).

Ecosystems and their constituent communities are highly dynamic in time (e.g., Brokaw 1985, Tscharntke et al. 2005, Malard et al. 2006, Bertrand et al. 2016), and temporal variation in community composition can impair the analysis of metacommunity diversity at a single date (Box 1). Temporal data should thus provide key information on community processes and assembly dynamics (Anderson and Cribble 1998, Magurran and Henderson 2010, Wolkowich et al. 2014, Buckley et al. 2018). To date, few studies have examined the temporal dynamics of metacommunities (Datry et al. 2016). They have mainly focused on describing spatiotemporal patterns (e.g. Soininen 2010, White et al. 2010, Legendre & Gauthier 2014). We here argue that such limited emphasis reflects (i) a lack of a general quantitative framework to analyse temporal changes (but see e.g., Nuvoloni et al. 2016) and (ii) the scarcity of proper empirical datasets.

#### Box 1

Fig. 1. Thought experiment demonstrating the benefits of a spatiotemporal analysis of community composition.

**Fig. 1.**
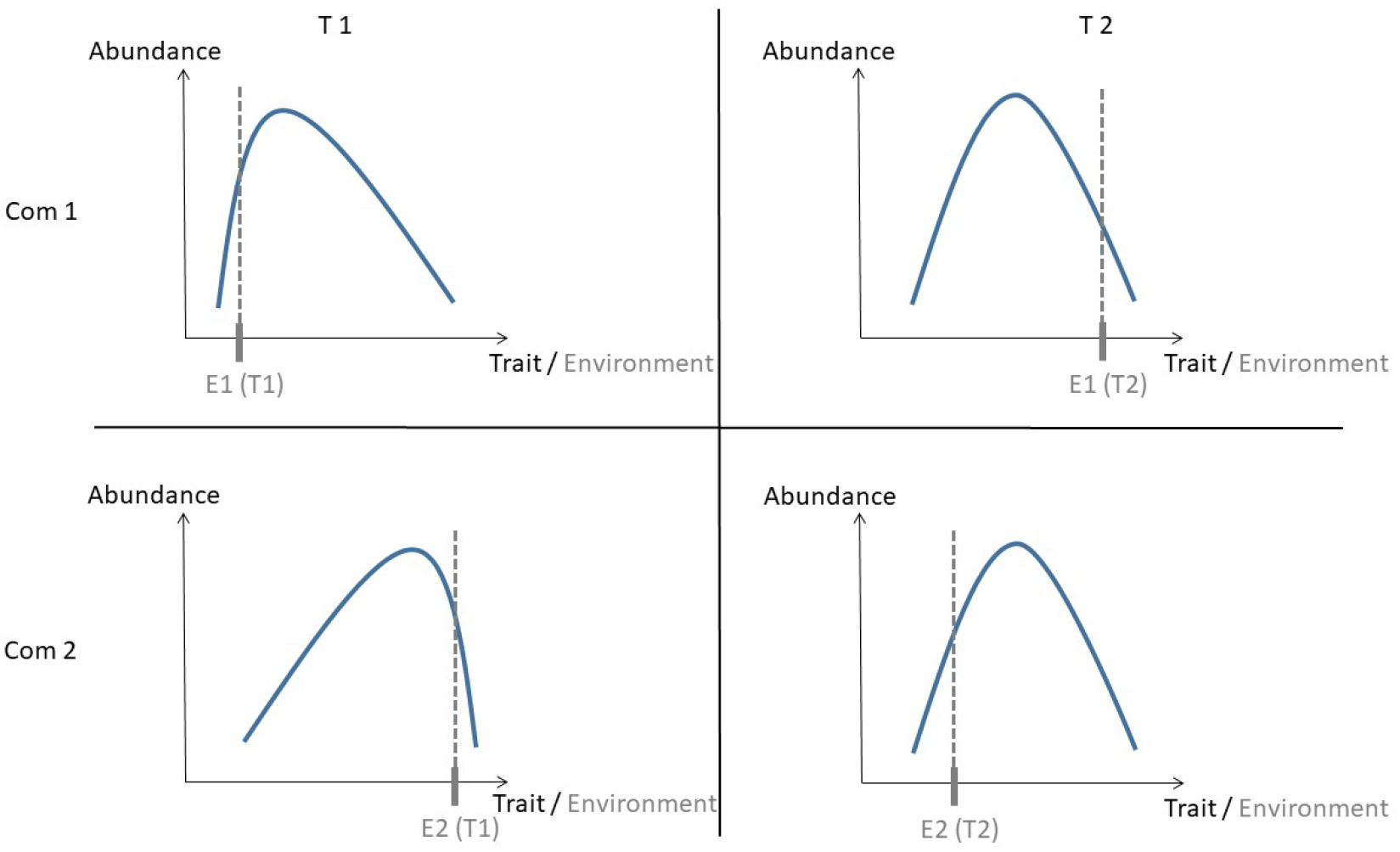
Thought experiment demonstrating the benefits of a spatiotemporal analysis of community composition. Blue curves represent the trait distribution of the community.

Imagine two communities with limited dispersal between each other and strongly structured by a one-dimensional environmental filter that selects species according to their (uni-dimensional) trait (Figure 1). At the first time step, community 1 experiences a low environmental value E_1_(t_1_) and is mainly composed of species with low trait values (Fig. 1a), while community 2 experiences a large environmental value E_2_(t_1_) and is mainly composed of species with large trait values (Fig. 1b). During the next time step, community 1 and 2 experiences large and low environmental values respectively, that enable some dispersers from the other community to settle in (Fig. 1c,d). This leads to a similar species composition of the two communities, despite the fact that current environmental conditions differ between the two communities. If one solely performs a spatial analysis of this metacommunity at time step 2, one would conclude that the environmental variable does not influence communities and thus that metacommunity dynamics is neutral with little dispersal limitation. If one instead looks at the temporal trajectories of the two communities (i.e. performs a spatiotemporal analysis), one would detect that community 1 and 2 moves to larger and lower trait values respectively, between t_1_ and t_2_, in a positively correlated manner with temporal environmental variations. One would also evidence that this tracking of environmental variation by community composition is only partial since abundant species do not have optimal trait values. This would provide evidence of some level of dispersal limitation between the two communities.

Nuvoloni et al. (2016) proposed to analyse the temporal turnover of community composition and to relate local turnover to environmental variables (see also Legendre 2019). We here propose to generalize this approach to metacommunities, that is to jointly analyse spatial and temporal turnovers of community composition. Indeed, using temporal signatures in metacommunity analyses is likely to reveal previously undetected environmental effects (Box 1). We therefore propose to perform path analyses that incorporate the influence of environmental, dispersal and community context on spatio-temporal turnovers of community composition. This approach presents the advantage of being sufficiently simple to elucidate the complex direct and indirect relationships among the drivers.

We propose a heuristic path model (Fig. 2). In this model, we predict that dispersal limitation and environmental filtering should entail positive correlations between community dissimilarity and, respectively, geographical distance and environmental distance (Borcard et al. 1992). Second, demographic stochasticity should entail negative correlations between mean community size and community dissimilarity across space and time, and positive correlation between temporal distance and community dissimilarity (Lande et al. 2003). Third, differences in community size should be positively linked to differences in species richness due to a general nestedness pattern of occurrence of abundant versus rare species (Srivastava & Lawton 1998), which in turn should entail greater community dissimilarity (through nestedness, see Baselga 2010). Finally, we consider that environmental distance can be correlated with temporal and geographical distances. Our heuristic understanding of spatio-temporal community dissimilarity patterns makes use of both direct and indirect relationships between explanatory variables that are themselves likely to be correlated to some extent. Path analyses therefore constitute a natural way to investigate such direct and indirect putative drivers of metacommunity dynamics (Kingsolver & Schemske 1991). Our heuristic path model is based on relationships between drivers and community composition that have been recurrently evidenced in the literature. Still, one may think of particular systems that may deviate from these general relationships and would require building of alternative heuristic path models.

**Fig. 2.**
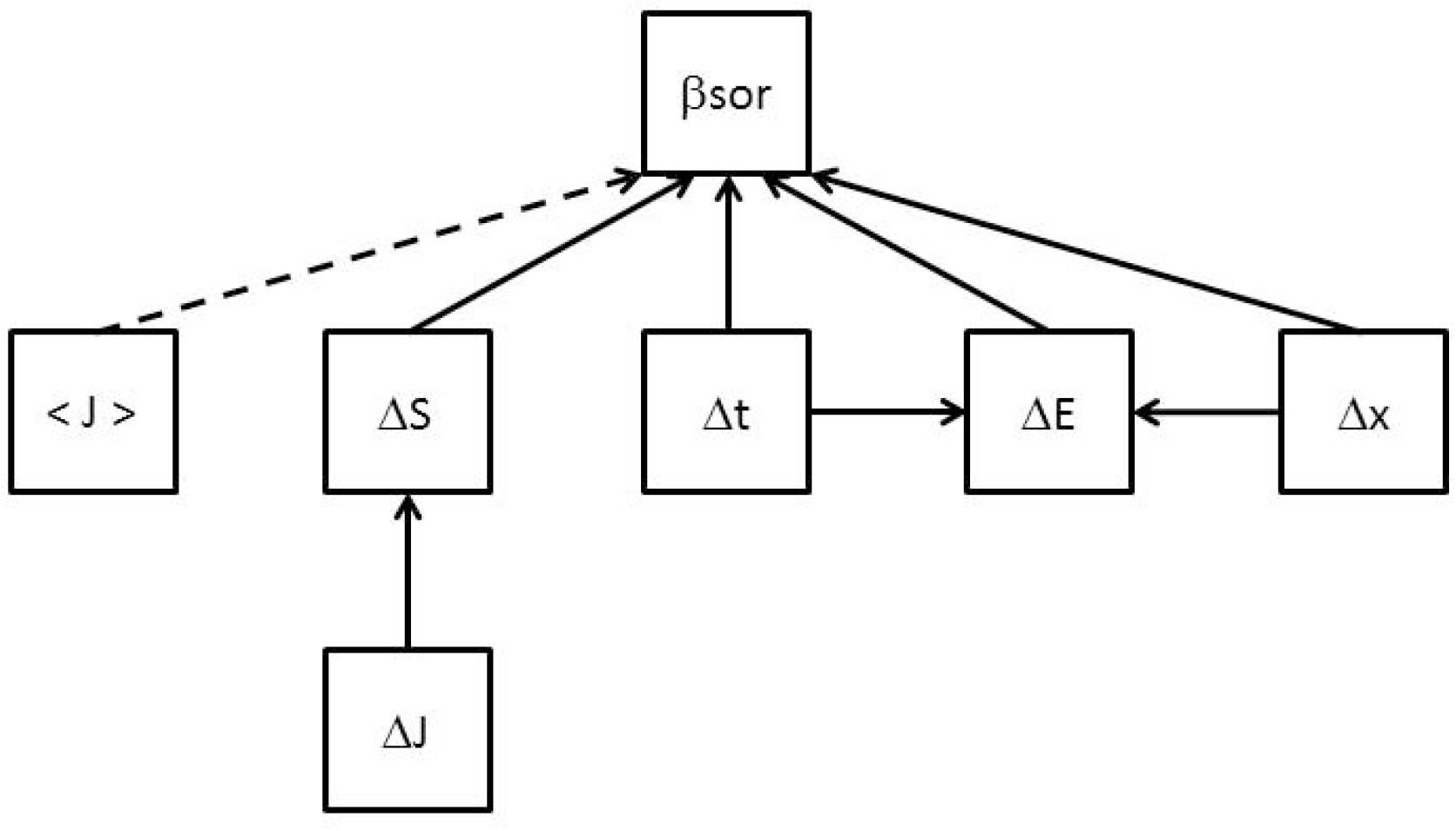
Heuristic path model to test the signature of ecological processes on spatiotemporal diversity patterns. A dashed or plain arrow represents a negative or positive correlation, respectively. <J> stands for the average community size in the metacommunity, t for time, x for space, E for the local environment and S for species richness. ∆ values represent difference of statistics in space and time. For instance, because it controls the intensity of ecological drift, the average community size is expected to negatively affect spatial and temporal diversity turnovers (negative arrow between <J> and β diversity).

We use this novel framework of spatio-temporal metacommunity analysis to analyse simulated data. We demonstrate that it enables us to detect the signature of the simulated metacommunity processes on the community patterns of beta-diversity through space and time. We then apply this framework to four real case studies. We find that spatial and temporal distances both influence dissimilarities in community composition. This effect is either direct or indirect through the spatio-temporal variations of the environmental drivers of community filtering.

## Materials and methods

### Assessing the path analysis framework with simulations

We devised an individual-based simulation algorithm of metacommunity dynamics in discrete time where communities are distributed across a two-dimensional grid. Our modelling choices were guided by the objectives of incorporating demographic stochasticity (using stochastic birth-death processes, Hubbell 2001), dispersal, environmental filtering (using the match between species traits and environmental values, Gravel et al. 2006), and environmental variability in space and time. This metacommunity model is particularly suited to represent sessile or territorial organisms, for which dispersal mostly occurs at the seed, larval or juvenile stages.

### Metacommunity simulator

#### Regional pool

We considered a fixed regional species pool of S species (S=100), each species i having a fixed regional frequency χ_i_ and a fixed trait value τ_i_, which corresponds to its environmental optimum. All species have the same regional frequency (χ_i_ =0.01) and trait values are regularly spaced between 0 and 1 (τ_i_ =i/100), so as to be parsimonious.

#### Landscape

We considered a gridded landscape of 400 cells (20 x 20) with fixed null boundary conditions. Abiotic environmental conditions within each cell at position (i,j) in the grid are assumed homogeneous and measured with the environmental variable E_ij_(t), which can vary in time (t). This variable influences processes of adult mortality and propagule establishment in each cell. There are J_ij_(t) individuals per cell, this number varying across space and time, depending on the balance between recruitment/immigration and mortality in each cell.

#### Environmental dynamics

The environmental variable E_ij_(t) in cell (i,j) at time t is decomposed into three components:

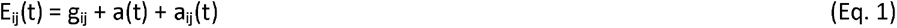

where g_ij_ represents an average environmental context in cell (i,j), a_t_ represents a temporal trend common to all cells, and a_ij_(t) represents a cell-specific temporal anomaly.

We consider a linear environmental gradient from the left to the right of the grid, so that g_ij_ regularly varies from 0.5 – e_1_/2 to 0.5 + e_1_/2 according to the column j of the cell (i,j), g_ij_ being constant on each column; a(t) is a triangle wave between - e_2_/2 and e_2_/2 with a period τ; a_ij_(t) is uniformly drawn between - e_3_/2 and e_3_/2 at each time step t and for each cell (i,j).

Environmental dynamics are parameterized with three parameters e_1_, e_2_ and e_3_ to control the magnitude of the spatial environmental gradient (e1), the temporal gradient (e2), and the residual environmental variability (e3). The triangle wave function is used so as to reach a dynamical equilibrium during the burn-in period, but a large wave period is used, so that a monotonic temporal gradient is actually simulated during the recorded dynamics.

#### Community dynamics

In each cell and during each time step, community dynamics is governed by four processes taking place sequentially: 1) reproduction, 2) propagule dispersal, 3) adult mortality and 4) propagule establishment. All cells are simultaneously updated.

##### 1) Reproduction

Each individual of the community produces propagules at a constant rate r so that the number of propagules produced by each individual during one time step is a Poisson draw with parameter r (with r ≤1).

##### 2) Dispersal

Two local dispersal models are alternatively used. In these two models, a proportion (1-m) of the propagules stays in their home cell, while a proportion m disperses outside the cell. In the global dispersal model, propagules homogeneously disperse among the 400 cells of the landscape, while in the neighbour dispersal model, they only disperse to the eight neighbouring cells (uniform random draws). This neighbour dispersal process enables us to model the dispersal influence on the spatial auto-correlation of species abundances. On top of this local dispersal, additional propagules arrive from the regional pool at a constant rate I in each cell, so that the number of long-distance dispersal propagules is computed as a Poisson draw with parameter I. This spatially-implicit long-distance dispersal process enables us to maintain species diversity at the grid scale (Hubbell 2001). This simulation setting leads to realistic uneven species-abundance distributions at the grid scale, despite the even species abundance distribution in the regional pool.

##### 3) Mortality

Each individual of species s has a local fitness f_s_(i,j,t) in cell (i,j) at time t, depending on the match between its trait value τ_s_ and the environmental variable E_ij_(t) in cell (i,j) at time t:

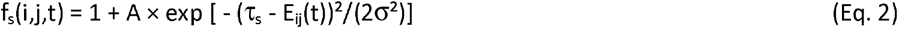

where parameter A controls the strength of environmental filtering (complete maladaptation leads to a local fitness of 1 while perfect adaptation to a local fitness of 1 + A) and parameter σ controls its specificity (a relatively good local adaptation is obtained when |τ_s_ - E_ij_(t)| is less than σ). This Gaussian-based equation is a rather standard way of modelling environmental filtering in community ecology (Gravel et al. 2006, Gilbert & Bennett 2010, Jabot 2010, Münkemüller & Gallien 2015, Sokol et al. 2017, Munoz et al. 2018)

The survival of adult individuals of species s is modelled at each time step t in cell (i,j) as a Bernoulli draw with probability (1-r) × f_s_(i,j,t) / (1+A). It implies that the individual death probability is at least equal to r and increases as individual fitness decreases.

##### 4) Establishment

Each cell has a carrying capacity of J individuals. The number of recruited individuals N_r_(i,j,t) in a cell (i,j) at time t follows a Poisson distribution with mean equal to J-N_ij_(t), where N_ij_(t) is the number of surviving adults in the cell after the mortality step. If N_ij_(t) is larger than J, then no individual is recruited. This modelling choice enables the number of individuals per cell to vary. The number of N_r_(i,j,t) recruited individuals follows a multinomial draw with probabilities proportional to species numbers of propagules reaching the focal cell times their local fitness f_s_(i,j,t).

The metacommunity is initialized with a multinomial draw of J=100 individuals from the regional pool in each cell. A burn-in period of 1,000 steps is used. Metacommunity dynamics then run for 20 time steps. 50 cells are randomly selected at the beginning of the simulation and their composition is recorded at each time step for subsequent analyses. The code of this simulator is provided (Supplementary material S1).

### Simulated scenarios

We devised 6 scenarios representing various combinations of ecological processes (Table 1). The first two scenarios implemented neutral dynamics, with either a homogeneous dispersal in the landscape or a dispersal to neighbouring cells to cause dispersal limitation. The third and fourth scenarios implemented environmental filtering along either a spatial or a temporal environmental gradient with homogeneous dispersal. The fifth and sixth scenarios implemented environmental filtering along simultaneous spatial and temporal gradients with either homogeneous dispersal or dispersal to neighbouring cells. Detailed parameter settings of the different scenarios are given in Table S2.

**Table 1.** Ecological processes and environmental spatiotemporal variations included in the six simulated scenarios.

| Scenario | Ecological processes |  |  | Environmental variables |  |
| --- | --- | --- | --- | --- | --- |
| | Global dispersal | Neighbour dispersal | Environmental filtering | Spatial ($e_1$) | Temporal ($e_2$) |
| 1 | + | - | - | - | - |
| 2 | - | + | - | - | - |
| 3 | + | - | + | + | - |
| 4 | + | - | + | - | + |
| 5 | + | - | + | + | + |
| 6 | - | + | + | + | + |

### Path analyses

We computed Sorensen dissimilarity indices for all pairs of sampled communities. Dissimilarity of communities at a given date represents spatial dissimilarity, dissimilarity of communities at the same site but at different dates represents temporal dissimilarity, and dissimilarity of the remaining pairs represents spatio-temporal dissimilarity. We computed spatial distances (Δx), temporal distances (Δt) and environmental distances (ΔE) for each pair of communities, as well as their mean community size (<J>), their absolute difference in community size (ΔJ) and in species richness (ΔS).

We ran a path analysis based on the heuristic causal model (Fig. 2) with the function “sem” of the R package “lavaan” (Rosseel 2012). Since path analyses were based on distance matrices, we used the permutation-based approach developed by Fourtune et al. (2018) taking into account non-independence and allowing to test for the significance of each path. We followed a Benjamini-Hochberg procedure to adjust the significance criterion (of 1%) for multiple testing. We assessed model fit with the Standardized Root Mean Square Residual (SRMR).

### Empirical datasets

We applied our conceptual framework to four empirical case studies.

#### Freshwater fishes

We analysed yearly samples of freshwater fish communities from the French Office for Biodiversity database including more than 1500 sites in France (Poulet et al. 2011). We analysed the dynamics of a subset metacommunity in the Garonne-Dordogne river drainage (Fourtune et al. 2016), including 32 sites monitored each year between 1995 and 2011, and for which precise environmental variables were available. This dataset included 51 fish species, for a total of 257,393 sampled individuals. Six environmental variables were recorded for each site: elevation, slope, average temperature in January 2011, average temperature in July 2011, width of the minor riverbed, and width of the water slide. The first five variables were temporally constant, while the last variable varied from year to year. Geographical distance between sites was computed along the river using the Carthage dataset of the French National Geographical Institute. We used log-transformed distances in analyses below, but results proved qualitatively similar when using raw distances.

#### Aquatic invertebrates

We compiled aquatic invertebrate communities across the Rhône river drainage in France. Benthic invertebrates were sampled on 6 sites of 11 different watersheds for a total of 66 sites. They were sampled for six months consecutively from the end of autumn to the beginning of summer for two years, 2014 and 2015, for a total of 12 sampling dates. The rivers considered are intermittent; when some sites were dry, they were not sampled at this date. Invertebrates were identified to the genus level but information was kept at the family level when no taxa were identified at the genus level for this family, resulting in a total of 231 taxa. Five environmental variables were measured at each site and sampling date: temperature, pH, conductivity, concentration in dioxygen and number of days since the last rewetting event of the watershed. We also computed log-transformed Euclidean geographic distances between sites.

#### Freshwater snails

The third dataset concerns the malacological fauna – 27 species - of a freshwater ponds network in the Guadeloupe Island. 250 sites have been yearly sampled since 2001 (17 years), with species density information. Species densities were multiplied by pond area to obtain estimated species abundances in each pond, and were then log-transformed. Each site is characterized by six temporally constant environmental variables (size, depth, vegetation cover, water quality, litter and a synthetic index of hydrological and vegetation stability, see Lamy et al. 2013 for additional details), and one temporally varying but spatially constant variable (annual rainfall). We calculated log-transformed, Euclidean geographical distances among sites. Missing data and empty sites were removed prior to analyses leading to a total of ca. 2800 samples.

#### Aquatic plants

We compiled aquatic plant communities in shallow lakes used for fish farming. These lakes are generally dried out during one year every three years. Twenty-four lakes were sampled from 2 to 7 years between 2008 and 2015, for a total of 81 sampling events and 84 sampled species (Arthaud et al. 2013). Average species cover was multiplied by lake areas to obtain estimated species abundances. Two environmental variables were used: chlorophyll a concentration reflecting water turbidity and light transmission, and the number of years since the last drying event.

## Results

### Analysis of simulated data

Our application of a causal modelling framework to simulated data validated our heuristic predictions and showed that the modelling framework allows reliable inferences of the ecological processes driving spatiotemporal variation in community composition for contrasted simulation scenarios. The effect of demographic stochasticity was detected in the four first scenarios with the moderate negative correlation between average community size and community dissimilarity (Fig. 3a-d). It was not detected in the last two scenarios, highlighting its minor importance compared to filtering processes in these scenarios. Demographic stochasticity was also detected with the moderate positive correlation between temporal distance and community dissimilarity in the two neutral scenarios (Fig. 3a-b). The effect of environmental filtering was detected in the four last scenarios with the positive correlation between environmental distance and community dissimilarity (Fig. 3c-f). The path analysis also correctly detected the positive correlation between spatial or temporal distances and environmental distance in the scenarios with a spatial or temporal environmental gradient respectively (Fig. 3c-f). In the scenarios with a temporal environmental gradient, a positive correlation between temporal distance and community dissimilarity was also identified (Fig. 3d-f). In these cases, we interpret this path as a signature of the inertia of these communities to the directional temporal environmental change simulated, rather than to a signature of demographic stochasticity. Indeed, the correlations were stronger than in the neutral scenarios in these cases. This result points that the interpretation of this specific path between temporal distance and community dissimilarity should be interpreted with caution, either as a signature of demographic stochasticity if temporal distance does not impact environmental distance, or as a signature of community inertia if temporal distance does impact environmental distance. In the two scenarios with dispersal limitation, a positive correlation between geographical distance and community dissimilarity was identified (Fig. 3b,f). A sampling effect was also detected in some scenarios with the positive correlations between differences in community size and differences in community richness and in turn between differences in community richness and community dissimilarity. These sampling effects were moderate when present and did not blur the other ecologically more informative paths. Finally, one can note the weak negative correlation between geographical distance and community dissimilarity in the third scenario with a spatial environmental gradient (Fig. 3c). We consider this correlation as an artefact due to the linear modelling framework used. It is outweighed by the strong positive indirect effect of geographical distance on community dissimilarity through environmental distance (Fig. 3c).

**Fig. 3.**
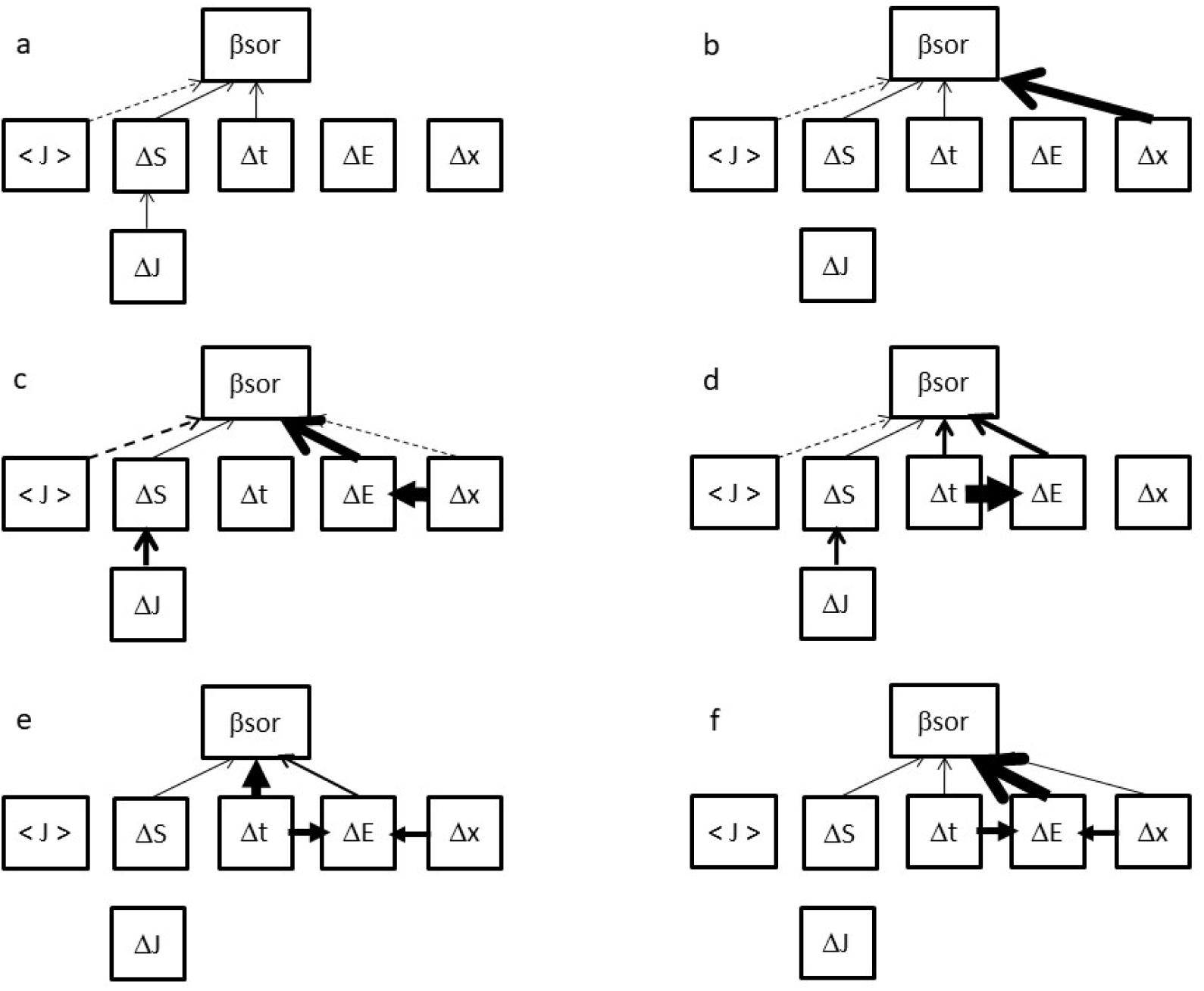
Path analyses on the six simulated scenarios. Arrows depict significant effects. Arrow width represents the strength of the standardized estimates. Numerical values are reported in Table S3.

### Analysis of empirical datasets

Our statistical framework revealed very consistent patterns across case studies (Fig. 4). The influence of demographic stochasticity was evidenced in all cases (see the dashed lines from <J> to βsor). Geographic distances Δx affected community dissimilarity (βsor) in all case studies, both directly (putatively through dispersal limitation) and indirectly through environmental distances ΔE. Environmental distances ΔE influenced community dissimilarity (βsor) in all case studies. Temporal distances Δt cascaded onto environmental distances in three of the four case studies. It also directly affected community dissimilarity in half of the case studies. Finally, differences in local species richness ΔS affected community dissimilarity in all case studies, and differences in local community sizes ΔJ influenced ΔS in three of the four case studies. The later result underlined that ΔS should be taken into account when assessing the drivers of community dissimilarity.

**Fig. 4.**
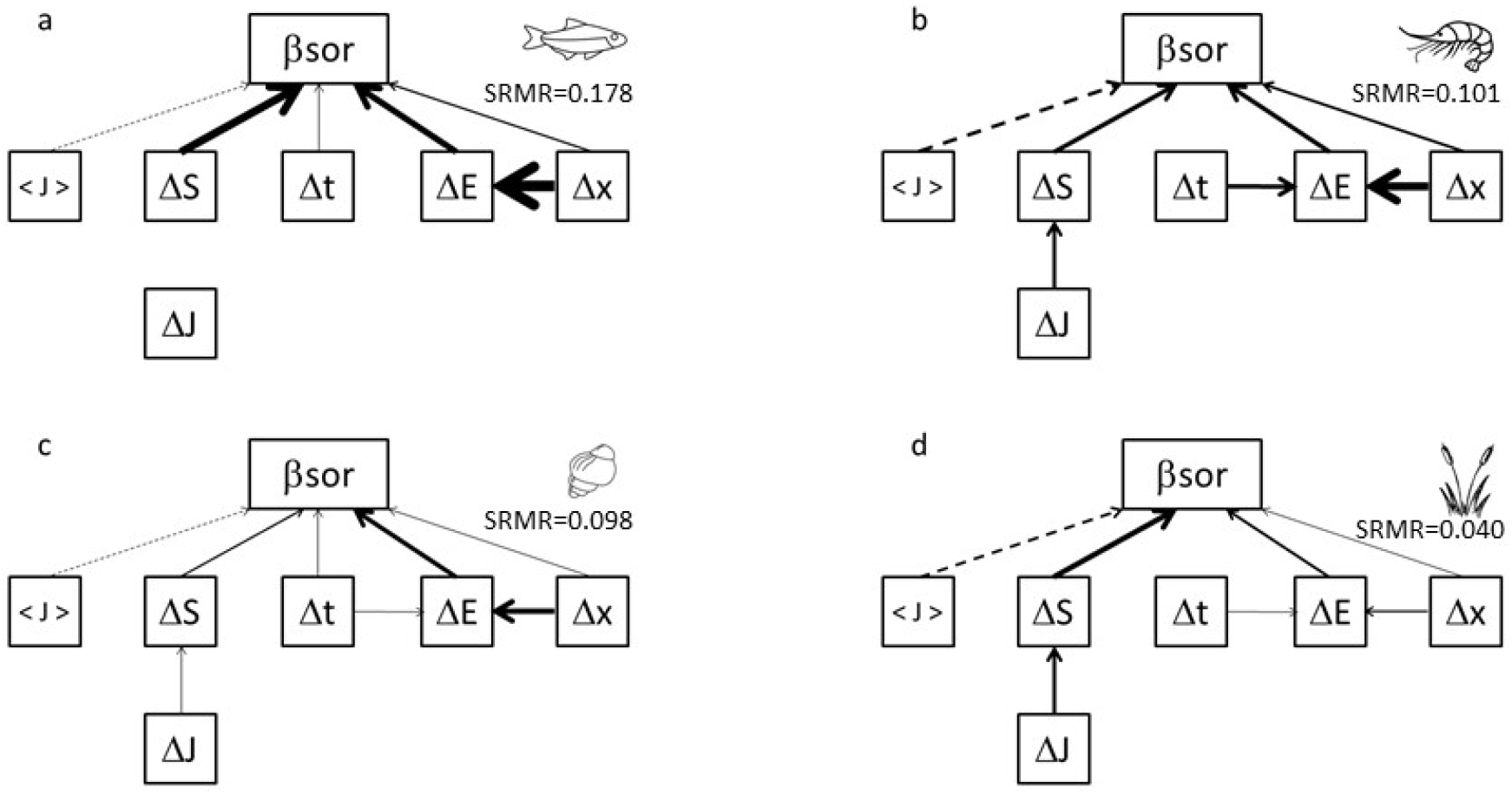
Path analyses for the four empirical datasets. a: freshwater fishes. b: aquatic invertebrates. c: molluscs. d: aquatic plants. Arrow width represents the strength of the standardized estimates. Dashed lines represent negative relationships. Paths from and towards ΔE were pooled in single arrows by summing the absolute values of the significant paths associated to each environmental variable. Only significant paths are shown. Numerical values of the standardized coefficients and of the associated p-values are reported in Tables S10-13. Values of the Standardized Root Mean Square Residual (SRMR) are mentioned for each dataset. Fish by Vladimir Belochkin, shrimp by Ana María Lora Macias, snail by Vega Asensio and cattail by Alex Muravev from the Noun Project.

Although we found support for the three main types of ecological drivers (demographic stochasticity, environmental variation and dispersal limitation), environmental variation was generally the strongest driver of community dissimilarity. This environmental variation was both spatially and temporally structured in three out of the four case studies (see the arrows from Δx and Δt towards ΔE). This result supports our call for an integrated spatiotemporal appraisal of metacommunity patterns.

## Discussion

### An operational approach to analyse spatiotemporal community turnover

We here proposed a simple approach to analyse spatiotemporal community turnover (Fig. 2). We found that approach allows detecting how dispersal, demographic stochasticity and environmental filtering influence metacommunity dynamics (Fig. 3). The proposed framework is robust and general since we examined strongly contrasted scenarios that all lead to path analysis results that were consistent with simulation choices. Applied to four real ecological case studies, we evidenced that the environmental drivers of community composition are always spatially auto-correlated and are temporally auto-correlated in three out of the four case studies. This further supports our call for a joint analysis of community turnover in both space and time.

### Detecting the contributions of entangled ecological processes

Applied to the fish data, this spatiotemporal framework revealed that the turnover in fish community composition at yearly and regional scales is mainly driven by environmental filtering (Fig. 4a) and that the environmental drivers are spatially auto-correlated (Δx ->ΔE). We also evidenced the influence of demographic stochasticity (ecological drift) and dispersal limitation on the spatiotemporal turnover component (Fig. 4a). Another main driver of community turnover is the heterogeneity in richness among local communities (ΔS), which we interpret as a nuisance variable here, since we do not have specific hypotheses on what may drive this heterogeneity beyond differences in community size (ΔJ). Alternative – yet non-exclusive – explanations for the observed variability in local species richness include the presence of a natural upstream-downstream gradient in species richness with more species near the outlet of the river networks (Muneepeerakul et al. 2008, Blanchet et al. 2014) and the introduction of non-native species that may not be homogeneous across the river network. Our analysis reveals that such potential drivers may have a dominant effect on the overall fish metacommunity structure at the regional scale.

Applied to the invertebrate data, the main driver of community turnover was also the heterogeneity in richness among local communities (Fig. 4b). This may result from the fact that this dataset comprises perennial and intermittent sites, and the latter ones generally harbour species-poor, original communities with taxa especially adapted to recover from disturbances (Datry et al. 2014). The other main drivers were demographic stochasticity and dispersal limitation, which may be explained by the intensity of local disturbances and regional fragmentation induced by drying events. Temporal and spatial distances also have a strong effect on environmental distances, as expected for intermittent rivers, as the stochasticity of drying events leads to a high spatiotemporal variability of the environment in a spatially and temporally auto-correlated way.

For the snail dataset, environmental variation was the main driver of community dissimilarity. Structuring environmental variables were vegetation and depth, reflecting a gradient from large, deep sites, with aquatic vegetation only near the shore, to small, shallow sites often entirely covered with vegetation. Shallow sites favour some pulmonate species that live near the surface, while most operculate snails are bottom-dwellers less dependent on macrophytes. Most environmental variables were spatially structured. Besides, communities were temporally structured. This may reflect a mix of long-term changes (invasion of new species and decline of some resident species during the study period) and short-term stochastic extinction-colonisation dynamics that make snail communities drift away from their initial states over time, within each site. For this last process, the strong negative impact of community size on species turnover indicates that small communities undergo faster temporal changes. These results point both to a strong impact of environmental filtering and fast temporal turnover, especially in small communities, which is in line with our experience with the ecology of freshwater snails.

For the aquatic plants dataset, our results point that environmental variation is a major driver of community dissimilarity, in accordance with the literature (Arthaud et al. 2013). Community size is also a significant driver of community composition. Indeed, in shallow lakes, communities are mostly made of plant species that are generally abundant (competitive or ruderal species, Arthaud et al. 2012), while a few species are characterized by lower population sizes (stress tolerant species). Our method evidenced that environmental filtering is both varying across space and time (Table S13). The light stress due to phytoplankton (chlorophyll a concentration) is temporally structured, because primary production varies strongly according to temperature and sunshine of the year. In contrast, the number of years since the last drying event is spatially structured, because of the specificities of the management performed at the catchment scale.

More generally, we found very consistent results across the four case studies despite the diversity of taxonomic groups (plants, aquatic invertebrates, molluscs and fishes) and habitats (lakes, ponds, perennial streams and intermittent rivers). It suggests the generality of the significance of spatiotemporal variation of environmental conditions for metacommunity dynamics. Ecologists should embrace a more dynamical view of metacommunity assembly and look beyond the predominant perspective, which considers communities as assembled through temporally fixed environmental filters. This present contribution offers a pragmatic way forward in this direction.

### Applying the proposed framework to metacommunity data

The proposed framework requires temporal data of metacommunity composition and temporal environmental variables that are thought to be influential for the system studied. If these environmental data are only available at the metacommunity scale, a different path model may still be used without the arrow between Δx and ΔE. Since the approach is exploratory, it does not require a minimal amount of sampled dates nor of sampled locations (beyond 2) to be operational. In the studied datasets, the number of sampled dates varied from 2 to 17, while the number of sampled locations varied from 24 to 250. Our approach relies on the analysis of community dissimilarity indices, so that it can be applied to species-rich communities that contain a substantial amount of rare species with low occurrence frequencies. The proposed approach is easy to conduct, since it does not require any advanced statistical training. It allows performing a first exploratory analysis of empirical data to assess the respective influences of complementary drivers of metacommunity dynamics (Kingsolver & Schemske 1991, Shipley 2000).

Some ecological systems may deviate from our general predictions for a variety of reasons. Users should then build alternative, biologically more relevant heuristic path models. Alternative path models can include different set of paths between the variables of Fig. 2, they can incorporate different signs for the predicted relationships, or they can incorporate other variables. For instance, environmental variables may display cyclic temporal dynamics. In such cases, it may be more pertinent to consider phase difference rather than absolute time difference (Δt). Similarly, the implicit assumption of spatial isotropy in the path model that makes use of geographical distances Δx may not be adapted to ecological systems in which the connectivity between sites is not solely driven by geographical distances (e.g., McRae et al. 2008). Another example is the one of disease or population outbreaks that travel through space and sometimes constitute a genuine environmental perturbation for entire communities (e.g., Tenow et al. 2013). In this case too, absolute time may not be a pertinent variable and may be fruitfully replaced by the state of outbreak (x-vt) where v is the speed of the travelling wave and x is the position of the site considered. More generally, when analysing transient systems, one should pay particular attention to the definition of paths involving time and to their interpretation. Our proposition is a simple and versatile approach to analyse standardized path coefficients, although this may not always be the choice to be favoured (Grace & Bollen 2005), so that researchers should evaluate the pros and cons of this choice for their particular case study.

Although the proposed framework appears powerful and robust, it is important to keep in mind that only simple linear relationships are modelled in the path analysis. Our analysis of simulated datasets supports this simple assumption (Fig. S4-90) and variable transformation procedures can be used to correct obvious non-linear relationships, as done here for some empirical case studies using log-transformation of geographical distances. Still, results should be solely interpreted as rough estimates of the respective influences of dispersal, demographic stochasticity and environmental filtering on community dynamics. Explored path models are not meant to be predictive. For such an endeavour, process-based dynamical models of metacommunity dynamics may be a much more suitable way forward (Evans et al. 2013, Mouquet et al. 2015). Such process-based dynamical models, however, require much more data on the system studied to be relevant. By identifying important drivers of metacommunity dynamics, the proposed framework can help design relevant process-based models that focus on the most influential processes.

The causal inference framework that we propose is a flexible way to make sense of temporal metacommunity data. We demonstrated its ability to detect influential ecological processes on simulated data. We illustrated its use on four real ecological datasets and explained how results can be interpreted. We finally explained how to deal with peculiarities of specific ecological systems within this framework, by modifying the path model. Beyond this methodological advance, our analyses of four sharply distinct ecological communities confirmed that environmental filtering is omnipresent and revealed that the environmental drivers of community composition do vary in both space and time, so that static metacommunity analyses should be abandoned in favour of a joint analysis of community turnover in both space and time.

## Acknowledgments

This work has been funded by the Irstea-Carnot project “MetaRISC”. This research was financed by the French government IDEX-ISITE initiative 16-IDEX-0001 (CAP 20-25). It has been partly undertaken at SETE, a laboratory which is part of the “Laboratoire d’Excellence” (LABEX) entitled TULIP (ANR-10-LABX-41). FM is funded by the CNRS and the ANR-funded ARSENIC and NGB projects (grants no. ANR-14-CE02-0012 and ANR-17-CE32-0011).

