## Supplementary Information for "Assessing metacommunity processes through signatures in spatiotemporal turnover of community composition"

S1: The C++ code to simulate the spatiotemporal dynamics of metacommunities is available at https://github.com/franckjabot/metacommunity_simulator; the R scripts to perform the path analysis of metacommunity spatio-temporal data and a tutorial for data formatting are available at https://github.com/franckjabot/metacommunity-analysis-script

Appendix S2: Additional information on the simulated scenarios.

Appendix S3: Additional information on empirical analyses.

### Data accessibility

Simulated datasets are available online:
https://doi.org/10.6084/m9.figshare.12090612.v1

The other empirical datasets are available here:
http://doi.org/10.5281/zenodo.3381338 (freshwater fishes dataset).
http://doi.org/10.5281/zenodo.3377490 (aquatic invertebrates dataset)
http://doi.org/10.5281/zenodo.3379615 (molluscs dataset)
http://doi.org/10.5281/zenodo.3383940 (aquatic plants dataset)

**Appendix S2: Additional information on the simulated scenarios.**

We here provide some details on the simulated scenarios.

| Scenario | I | m | A | $\sigma$ | $e_1$ | $e_2$ | $e_3$ | r | Glob. disp. |
|---|---|---|---|---|---|---|---|---|---|
| 1 | 0.1 | 1 | 0 | - | 0 | 0 | 0.1 | 0.5 | Y |
| 2 | 0.1 | 1 | 0 | - | 0 | 0 | 0.1 | 0.5 | N |
| 3 | 100 | 1 | 10 | 0.02 | 0.15 | 0 | 0.01 | 0.1 | Y |
| 4 | 100 | 1 | 10 | 0.02 | 0 | 0.5 | 0.01 | 0.1 | Y |
| 5 | 1 | 1 | 1 | 0.02 | 0.15 | 0.5 | 0.01 | 0.1 | Y |
| 6 | 1 | 1 | 1 | 0.02 | 0.15 | 0.5 | 0.01 | 0.1 | N |

**Table S2**: model parameters used in the simulated scenarios. Glob. disp. is a Boolean value indicating whether a global dispersal between all cells in the landscape was used (Y) or whether a local dispersal between neighbouring cells was used (N). In addition, all simulations were performed with a value of $\tau$ equal to 130.

| Scenario | 1 | 2 | 3 | 4 | 5 | 6 |
|---|---|---|---|---|---|---|
| $\beta_{sor} \leftarrow \langle J \rangle$ | **-0.07 – 0** | **-0.05 – 0.004** | **-0.14 – 0** | **-0.10 – 0** | -0.01 – 0.29 | -0.02 – 0.21 |
| $\beta_{sor} \leftarrow \Delta S$ | **0.10 – 0** | **0.07 – 0** | **0.04 – 0** | **0.04 – 0** | **0.04 – 0** | **0.05 – 0** |
| $\beta_{sor} \leftarrow \Delta t$ | **0.07 – 0** | **0.05 – 0** | -0.007 – 0.18 | **0.23 – 0** | **0.55 – 0** | **0.04 – 0** |
| $\beta_{sor} \leftarrow \Delta E$ | 0 – 0.48 | 0.001 – 0.43 | **0.55 – 0** | **0.33 – 0** | **0.14 – 0** | **0.64 – 0** |
| $\beta_{sor} \leftarrow \Delta x$ | 0.003 – 0.35 | **0.55 – 0** | **-0.03 – 0** | 0.001 – 0.44 | 0.009 – 0.16 | **0.07 – 0** |
| $\Delta S \leftarrow \Delta J$ | **0.05 – 0** | -0.002 – 0.50 | **0.27 – 0** | **0.20 – 0** | -0.007 – 0.32 | 0.01 – 0.21 |
| $\Delta E \leftarrow \Delta t$ | -0.008 – 0.09 | -0.01 – 0.06 | 0.001 – 0.46 | **0.99 – 0** | **0.41 – 0** | **0.39 – 0** |
| $\Delta E \leftarrow \Delta x$ | 0.005 – 0.21 | 0.003 – 0.32 | **0.74 – 0** | 0.002 – 0.38 | **0.28 – 0** | **0.33 – 0** |
| SRMR | 0.008 | 0.027 | 0.027 | 0.006 | 0.012 | 0.012 |

**Table S3**: Standardized estimates and p-values for the path analyses on simulated scenarios (Fig.4). p-values equal to 0 actually mean <0.001. Significant effects at the 1% level with a Benjamini-Hochberg correction are depicted in bold. The last line reports the Standardized Root Mean Square Residual (SRMR) that is a standard measure of model fit for path analyses.

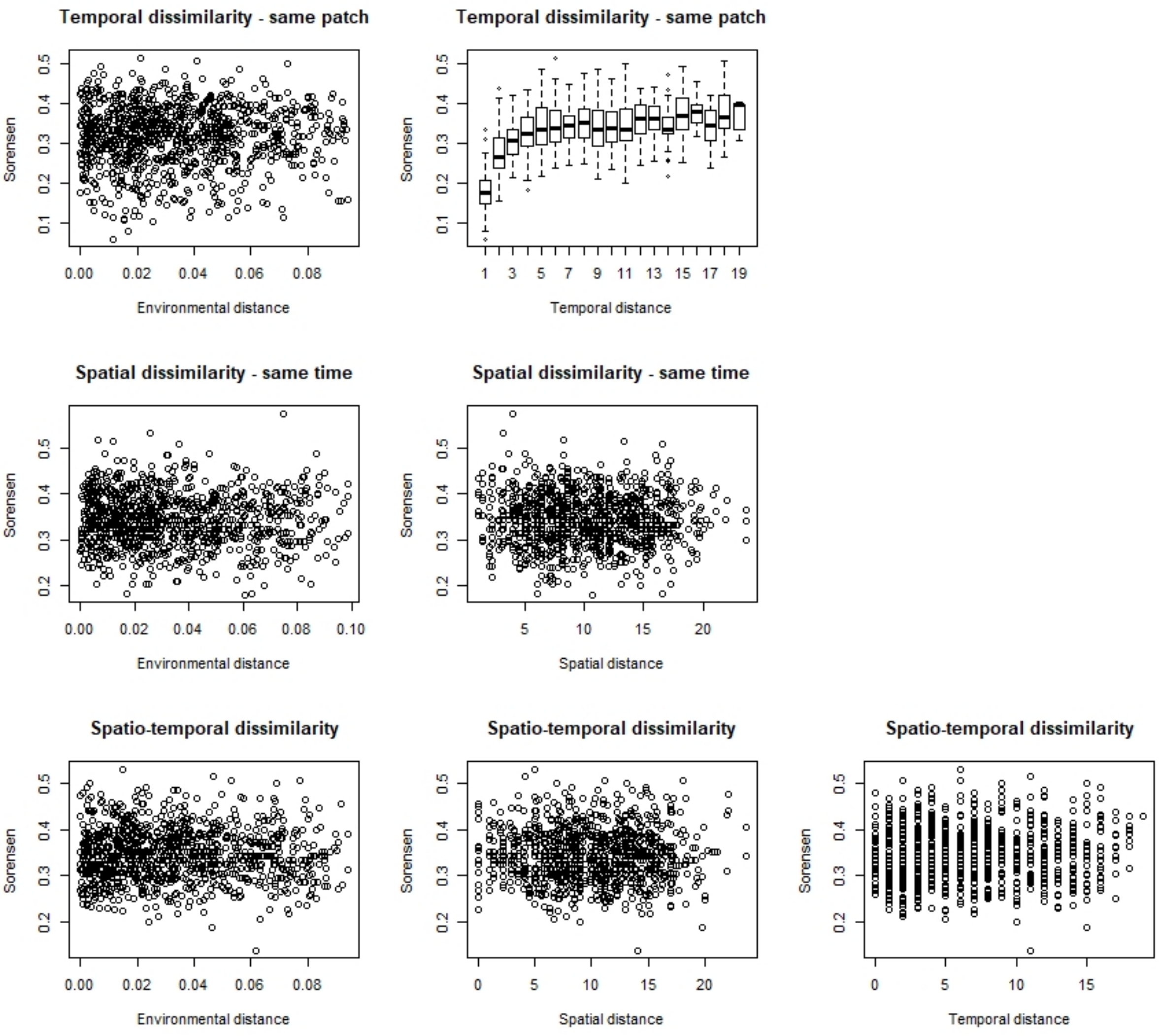


**Fig. S4** – Descriptive plots for the first scenario (neutral without dispersal limitation)

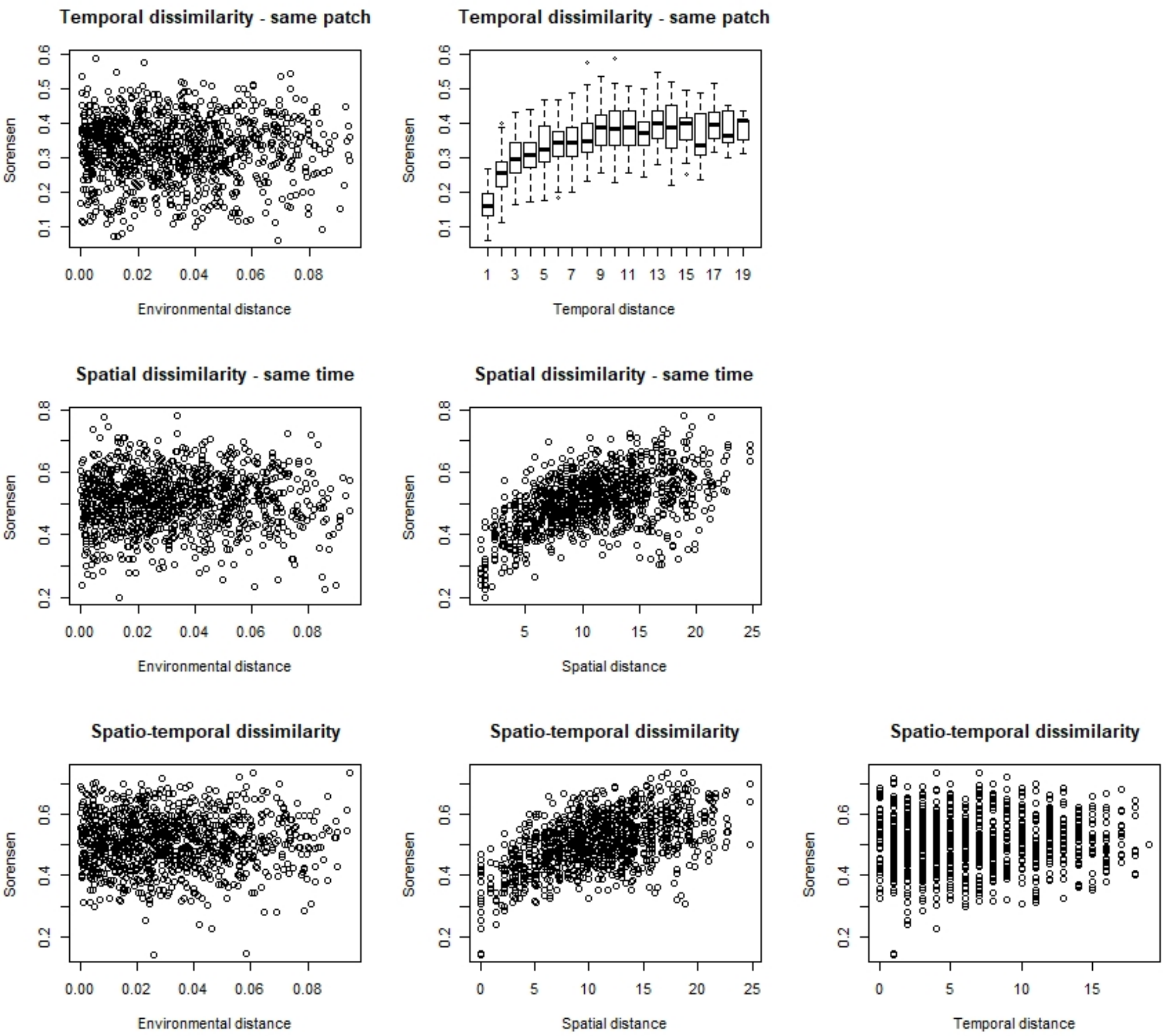


**Fig. S5** – Descriptive plots for the second scenario (neutral with dispersal limitation)

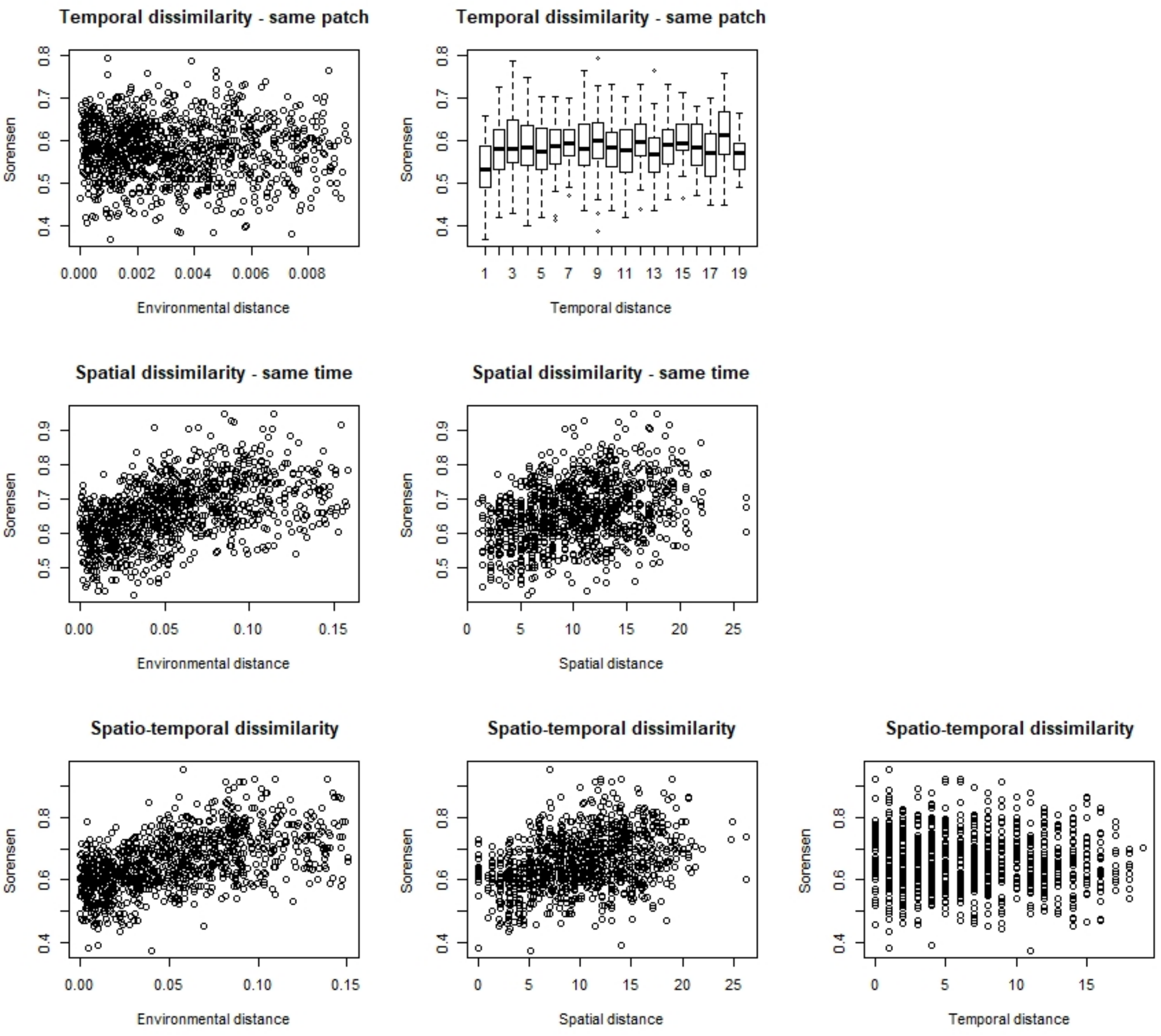


**Fig. S6** – Descriptive plots for the third scenario (spatial environmental gradient without dispersal limitation)

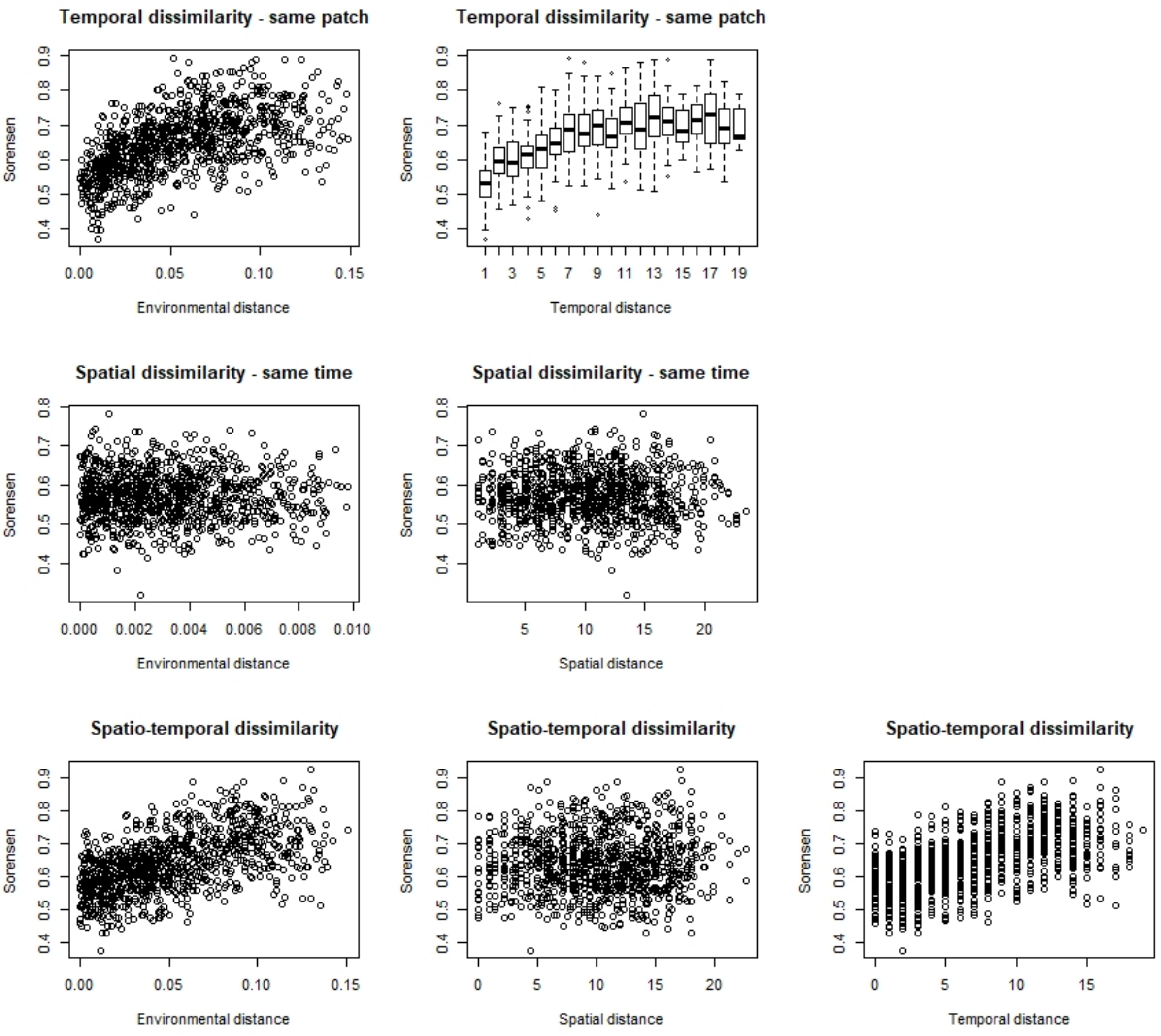


**Fig. S7** – Descriptive plots for the fourth scenario (temporal environmental gradient without dispersal limitation)

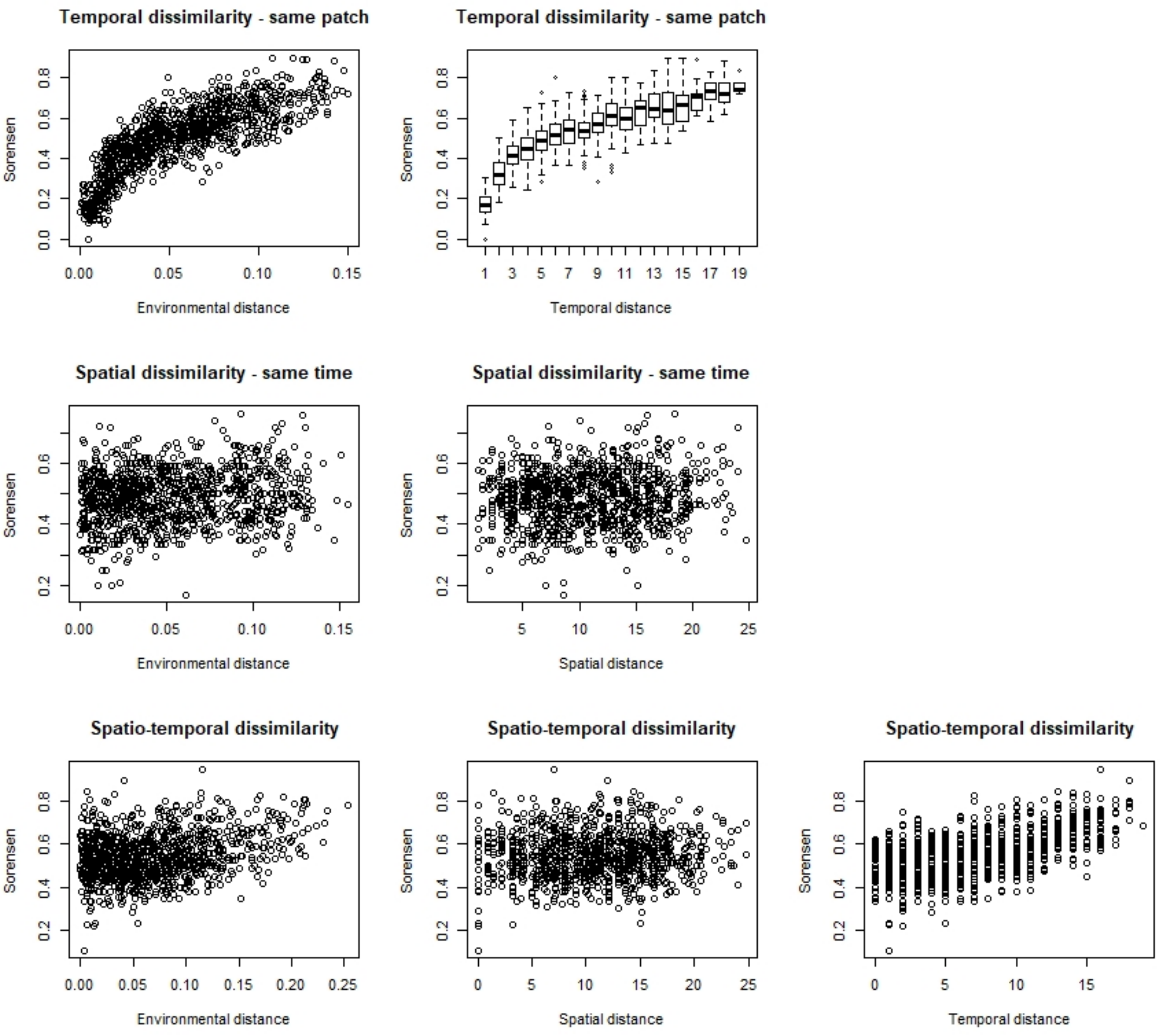


**Fig. S8** – Descriptive plots for the fifth scenario (spatial and temporal environmental gradients without dispersal limitation)

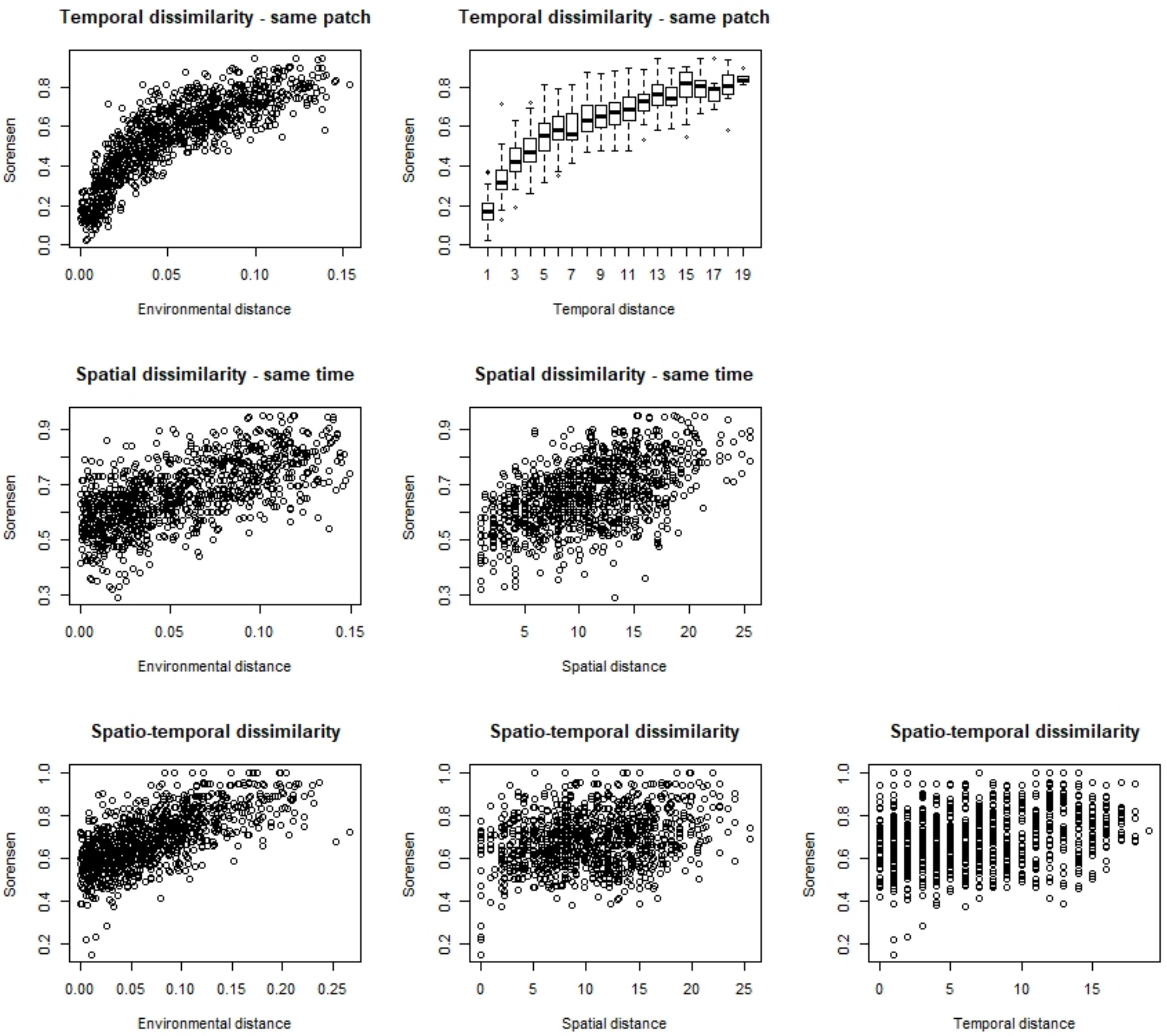


**Fig. S9** – Descriptive plots for the sixth scenario (spatial and temporal environmental gradients with dispersal limitation)

**Appendix S3: Additional information on empirical analyses.**

We here provide the numerical results of the path analyses conducted for the four datasets (Tables S10-13).

| Environmental variables | | Width of the water slide | Width of the minor bed | Elevation | Slope | Average temperature in January 2011 | Average temperature in July 2011 |
|---|---|---|---|---|---|---|---|
| $\beta_{sor} \leftarrow \langle J \rangle$ | **-0.06 – 0.001** | | | | | | |
| $\beta_{sor} \leftarrow \Delta S$ | **0.69 – 0** | | | | | | |
| $\beta_{sor} \leftarrow \Delta t$ | **0.02 – 0.02** | | | | | | |
| $\beta_{sor} \leftarrow \Delta E$ | | **0.04 – 0.002** | **0.08 – 0** | **0.19 – 0** | **0.13 – 0** | **0.04 – 0.004** | **0.03 – 0.03** |
| $\beta_{sor} \leftarrow \Delta x$ | **0.13 – 0** | | | | | | |
| $\Delta S \leftarrow \Delta J$ | 0.03 – 0.09 | | | | | | |
| $\Delta E \leftarrow \Delta t$ | | 0.007 – 0.36 | | | | | |
| $\Delta E \leftarrow \Delta x$ | | **0.06 – 0.01** | **0.07 – 0.007** | **0.25 – 0** | **0.17 – 0** | **0.26 – 0** | **0.12 – 0** |

**Table S10**: Standardized estimates and p-values for the path analysis of the AFB freshwater fish dataset. p-values equal to 0 actually mean <0.001. Significant effects at the 5% level with a Benjamini-Hochberg correction are depicted in bold. SRMR = 0.178.

| Environmental variables | | Temperature | pH | Conductivity | Concentration in dioxygen | Number of days since the last rewetting event of the watershed |
|---|---|---|---|---|---|---|
| $\beta_{sor} \leftarrow \langle J \rangle$ | **-0.31 – 0** | | | | | |
| $\beta_{sor} \leftarrow \Delta S$ | **0.47 – 0** | | | | | |
| $\beta_{sor} \leftarrow \Delta t$ | 0.01 – 0.17 | | | | | |
| $\beta_{sor} \leftarrow \Delta E$ | | **0.07 – 0** | **0.11 – 0** | **0.10 – 0** | **-0.06 – 0** | **0.03 – 0.02** |
| $\beta_{sor} \leftarrow \Delta x$ | **0.25 – 0** | | | | | |
| $\Delta S \leftarrow \Delta J$ | **0.28 – 0** | | | | | |
| $\Delta E \leftarrow \Delta t$ | | **0.08 – 0** | **0.11 – 0** | **0.04 – 0** | 0.02 – 0.08 | **0.18 – 0** |
| $\Delta E \leftarrow \Delta x$ | | **0.02 – 0.01** | **0.27 – 0** | **0.31 – 0** | **0.04 – 0** | **0.02 – 0** |

**Table S11**: Standardized estimates and p-values for the path analysis of the Irstea aquatic invertebrate dataset. p-values equal to 0 actually mean <0.001. Significant effects at the 5% level with a Benjamini-Hochberg correction are depicted in bold.

| Environmental variables | | Pond size | Pond depth | Vegetation cover | Water quality | Litter amount | Stability | Annual rainfall |
|---|---|---|---|---|---|---|---|---|
| $\beta_{sor} \leftarrow \langle J \rangle$ | **-0.10 – 0** | | | | | | | |
| $\beta_{sor} \leftarrow \Delta S$ | **0.12 – 0** | | | | | | | |
| $\beta_{sor} \leftarrow \Delta t$ | **0.08 – 0** | | | | | | | |
| $\beta_{sor} \leftarrow \Delta E$ | | 0 – 0.32 | **0.10 – 0** | **0.15 – 0** | **0.06 – 0** | **-0.02 – 0.005** | **0.05 – 0** | 0.01 – 0.06 |
| $\beta_{sor} \leftarrow \Delta x$ | **0.06 – 0** | | | | | | | |
| $\Delta S \leftarrow \Delta J$ | **0.06 – 0** | | | | | | | |
| $\Delta E \leftarrow \Delta t$ | | | | | | | | **0.09 – 0** |
| $\Delta E \leftarrow \Delta x$ | | **0.07 – 0** | **0.02 – 0.004** | 0.01 – 0.10 | **0.07 – 0** | **0.13 – 0** | **0.15 – 0** | |

**Table S12**: Standardized estimates and p-values for the path analysis of the mollusc dataset. p-values equal to 0 actually mean <0.001. Significant effects at the 5% level with a Benjamini-Hochberg correction are depicted in bold.

| Environmental variables | | Chlorophyll a concentration | Number of years since the last drying event |
|---|---|---|---|
| $\beta_{sor} \leftarrow \langle J \rangle$ | **-0.24 – 0** | | |
| $\beta_{sor} \leftarrow \Delta S$ | **0.47 – 0** | | |
| $\beta_{sor} \leftarrow \Delta t$ | 0.01 – 0.33 | | |
| $\beta_{sor} \leftarrow \Delta E$ | | **0.16 – 0** | **-0.06 – 0.04** |
| $\beta_{sor} \leftarrow \Delta x$ | **0.09 – 0.01** | | |
| $\Delta S \leftarrow \Delta J$ | **0.29 – 0** | | |
| $\Delta E \leftarrow \Delta t$ | | **0.09 – 0.02** | -0.03 – 0.29 |
| $\Delta E \leftarrow \Delta x$ | | 0.002 – 0.44 | **0.13 – 0.001** |

**Table S13**: Standardized estimates and p-values for the path analysis of the aquatic plant dataset. p-values equal to 0 actually mean <0.001. Significant effects at the 5% level with a Benjamini-Hochberg correction are depicted in bold.
